# Discovery of Light-Powered Organic Ion Transport by a Natural Protein

**DOI:** 10.1101/2025.07.01.658226

**Authors:** Simiao Shen, Shoichiro Akita, Joji Wada, Mako Eguchi, Takashi Tsukamoto, Kwang-Hwan Jung, Yuki Sudo, Takashi Kikukawa

## Abstract

Membrane transport proteins play vital roles in living cells by selectively transporting ions and molecules across biological membranes. Among these, microbial rhodopsins are unique in their unparalleled capacity to harness light to drive ion translocation. This distinctive feature has led to their widespread use as optogenetic tools in neuroscience, physiology, and biomedical applications. While not all microbial rhodopsins function as ion transporters, many ion-translocating variants have been discovered since the identification of the first member—a light-driven H^+^ pump—in the 1970s. These proteins share a compact structure composed of only seven transmembrane helices and have long been thought to specialize exclusively in transporting small inorganic ions such as H^+^, Cl^−^, and Na^+^. Here, we show that several anion-pumping microbial rhodopsins can also transport organic anions. In particular, a rhodopsin from cyanobacteria is capable of transporting bulky organic anions, including those containing benzene rings, with molecular volumes up to ∼120 Å^3^—five times that of Cl^−^. These organic ions bind to the dark state and are translocated upon photoactivation, following a mechanism similar to that of inorganic anion transport. Mutational analysis indicates that both classes of substrates share a common binding site. Only anions with p*K*a values below 2 were transported, suggesting that a retained negative charge is essential for binding to the dark state—a prerequisite for transport. This study expands the known substrate repertoire of microbial rhodopsins and introduces new possibilities for optogenetic strategies based on light-driven delivery of bioactive organic molecules.

## INTRODUCTION

Membrane transport proteins are indispensable for cellular function, enabling the selective movement of ions and molecules across biological membranes. Among them, microbial rhodopsins stand out for their ability to harness light energy to drive ion transport ^1-4^. Since the discovery of the first light-driven H^+^ pump in the 1970s ^5-6^, many ion-translocating variants have been identified ^1-4^. Their light-activated nature has made them valuable model systems for studying membrane transport and, more recently, powerful tools for optical manipulation of cells ^7-8^. Channelrhodopsins, which function as light-gated ion channels, are the most widely used in optogenetics ^9-10^, while ion-pumping rhodopsins have also demonstrated strong potential ^8, 11^. These proteins now enable light-based control over diverse cellular processes, including the activation or inhibition of excitable cells, modulation of signaling pathways, and induction of programmed cell death ^8, 11-12^. To date, all characterized microbial rhodopsins have been shown to transport only small inorganic ions such as H^+^, Cl^−^, or Na^+^. These ions, while effective for electrical or pH modulation, lack the intrinsic biological activity of organic compounds. Expanding the substrate scope of microbial rhodopsins to include organic ions could significantly enhance their utility in optogenetics, enabling light-driven delivery of bioactive molecules. Here, we demonstrate for the first time that certain microbial rhodopsins can transport organic anions.

Microbial rhodopsins contain the chromophore retinal within the protein ^1-4^. Fig. 1A shows the structure of a Cl^−^-pumping rhodopsin (ClR) from the cyanobacterium *Synechocystis* sp. PCC 7509, referred to here as SyHR. Upon light absorption, retinal undergoes isomerization, triggering a conformational change in the protein. These structural changes proceed through a cyclic series of intermediates known as the photocycle, ultimately returning the protein to its ground state. Each rhodopsin performs a specific function during this cycle within its native host.

**Figure 1.**
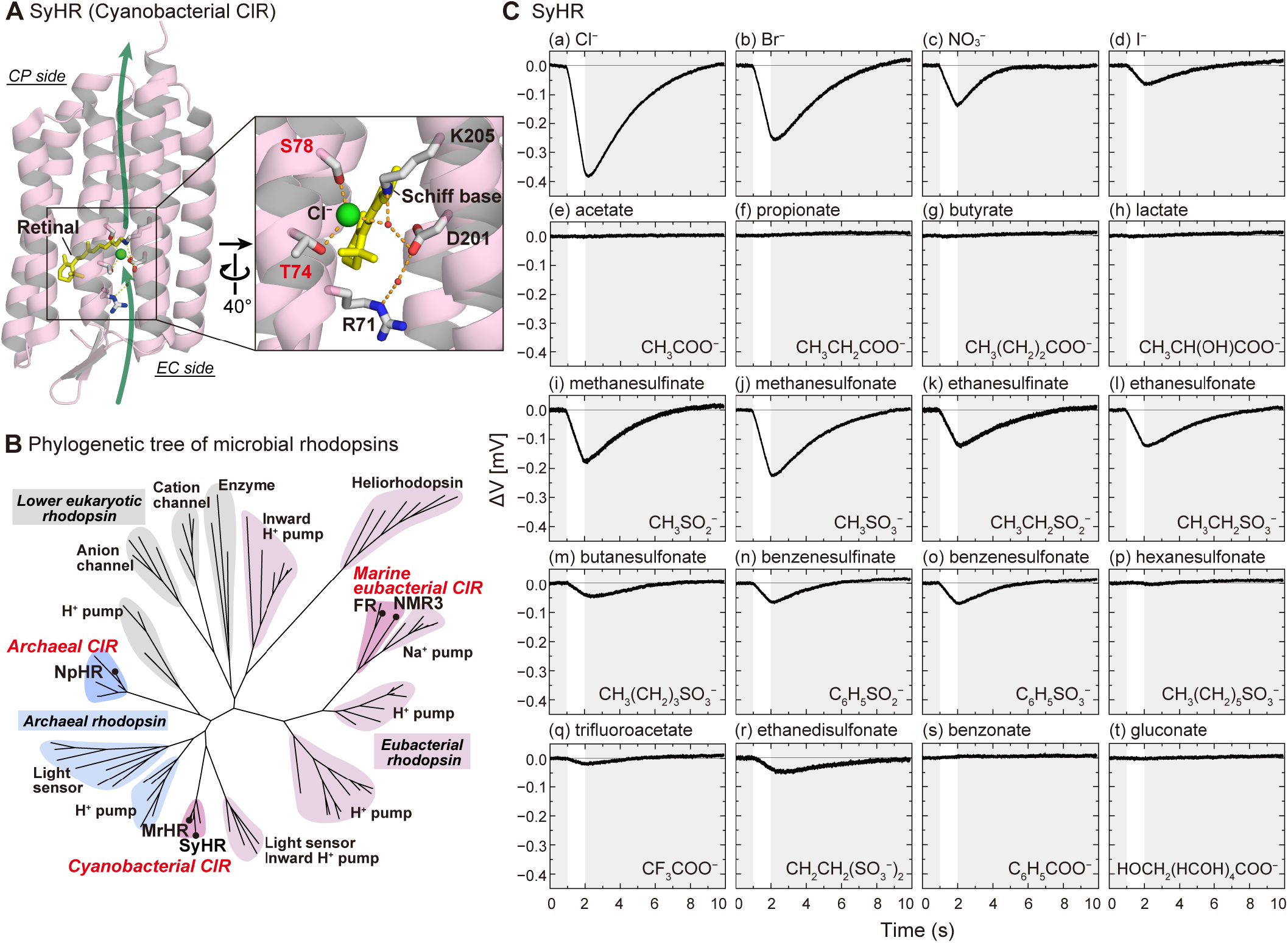
Tertiary structure, phylogenetic position, and anion transport activity of SyHR. (A) Tertiary structure of unphotolyzed SyHR in the Cl^−^-bound state (PDB ID: 7ZOU). Water molecules are shown as small spheres, and polar contacts within 3.2 Å are indicated by dashed lines. (B) Phylogenetic relationships among microbial rhodopsins. (C) Anion transport activity of SyHR. Each trace shows the light-induced voltage change recorded by the ITO electrode in the electrochemical cell. E. coli cells were suspended in 200 mM sodium salts of the respective anions with 10 μM CCCP. Actinic light was applied for 1–2 seconds.

Although many ion-translocating microbial rhodopsins have been identified over the past 50 years, they have been thought to transport only small inorganic ions—typically H^+^, Cl^−^, and Na^+^—an assumption consistent with their compact structure ^1-4^. While transporters for organic molecules typically consist of two homologous domains, each with 6–10 transmembrane helices, microbial rhodopsins comprise a single domain containing just seven. This structural simplicity has fostered the widespread belief that microbial rhodopsins are specialized for small inorganic ions and are incapable of transporting organic ions. In this study, we focused on ClRs, which were known to transport a few polyatomic inorganic anions, such as NO3^−^, SCN^−^, and SO_4_^2− 13-18^. Although these ions lack hydrocarbon groups, their volumes are close to those of organic anions like acetate and propionate—short-chain fatty acids (SCFAs) known for their bioactivity in animals. Contrary to expectations, we found that ClRs fail to transport SCFAs. However, several ClRs, and particularly SyHR, were capable of transporting other organic anions, including those with benzene rings.

To our knowledge, this is the first report demonstrating that naturally occurring proteins can use light energy to transport organic compounds across membranes.

## MATERIALS AND METHODS

### Protein expression, purification, and lipid reconstitution

*Escherichia coli* DH5α was used for DNA manipulation. Expression plasmids for SyHR, ClR from *Natronomonas pharaonis* (NpHR), and ClR from *Mastigocladopsis repens* (MrHR) were described previously and were based on the pET-21c vector ^16, 19-20^. The gene of ClR from *Fulvimarina pelagi* (FR) (NCBI accession ID: WP_038437030) was ampli-fied from the genomic DNA of *F. pelagi* strain DSM 15513 and inserted into the pKA001 vector under the control of the lacUV5 promoter ^21^. The gene of ClR from *Nonlabens marinus* S1-08 (NM-R3) (NCBI accession ID: BAO55276) was similarly inserted into the same site of pKA001 after codon optimization for *E. coli* expression. All constructs encoded proteins with a C-terminal hexahistidine tag. Point mutations in SyHR were introduced using the QuikChange Site-Directed Mutagenesis Kit (Agilent Technologies), and all DNA sequences were confirmed using standard procedures.

*E. coli* C43(DE3) was used to express all proteins. Expression followed previously described protocols ^22-23^. Briefly, cells were grown in 2×YT medium at 37 °C, and expression was induced by adding 1 mM isopropyl-β-D-thiogalactopyranoside and 10 μM all-*trans* retinal. After 3 h of induction, cells were harvested for downstream processing.

SyHR was purified and reconstituted into lipids as previously described ^24-25^. Briefly, harvested cells were lysed by sonication, and membrane fractions were collected and solubilized with 1.5% n-dodecyl β-D-maltopyranoside. Solubilized proteins were purified using nickel–nitrilotriacetic acid (Ni-NTA) agarose resin. Protein concentrations were determined based on absorbance at λ_max_ (∼538 nm) using an extinction coefficient of 39,100 M^−1^ cm^−1 17^. For lipid reconstitution, phosphatidylcholine (from egg yolk; Avanti) was added at a protein-to-lipid molar ratio of 1:50. Detergent was removed by gentle stirring overnight at 4 °C in the presence of SM2 Adsorbent BioBeads (Bio-Rad). Reconstituted proteins were collected by centrifugation and used in subsequent experiments.

### Anion pump activity measurements

. Suspensions of *E. coli* cells expressing ClR were prepared as described previously ^17, 26^. Briefly, harvested cells were washed twice with 0.2 M sodium salt solutions of the test anions, resuspended in the same solutions, and gently shaken overnight at 4 °C in the presence of 10 μM carbonyl cyanide mchlorophenylhydrazone (CCCP). On the following day, cells were washed again with the same solutions. Final cell density was adjusted to an optical density at 660 nm (OD_660_) of 10.0.

In the presence of CCCP, inward transport of anions by ClRs generates a negative membrane potential that passively drives H^+^ influx into the cells. This proton inflow leads to a light-induced pH increase in the suspension. To detect this, we used a custom-built photochemical cell (Fig. S1A) instead of a conventional pH electrode ^17^. The setup consisted of two indium tin oxide (ITO) electrodes separated by a dialysis membrane. One chamber contained the *E. coli* suspension; the other held a reference solution containing the same salt and 10 μM CCCP. The ITO electrodes are transparent and provide a rapid response to pH changes. Because the pH change occurs only in the cell suspension, a voltage difference arises between the electrodes, reflecting anion transport activity.

Anion pump activity was induced by a light from a 150 W xenon arc lamp filtered through two glass filters (IRA-25S and Y-50; Toshiba, Tokyo, Japan). Voltage changes were recorded for 10 s with illumination from 1–2 s. Each measurement was repeated 30–50 times, and average traces were used to improve the signal-to-noise ratio. The light intensity was 28.4 mW/cm^2^ at 530 nm, measured using an Orion-PD optical power meter (Ophir Optonics Ltd.). Measurements were conducted at room temperature (approximately 25 °C).

Anion transport activity was quantified as the slope (denoted α) of the light-induced voltage change. Slopes were calculated for each of 30–50 traces by linear fitting, and mean (⟨α⟩) and standard deviation values were computed.

Initial pH values of the suspensions were above 5.9 and primarily determined by the intrinsic buffer capacity of the *E. coli* cells. However, anions with relatively high p*K*a values (e.g., SCFAs) could act as buffers themselves and reduce the magnitude of the observed voltage change. Therefore, we measured the buffer capacity (β) of each sample and used this to normalize the α values. Specifically, 2 mL of each sample was placed in a cuvette, and pH was measured before and after the addition of 50 nmol of H^+^ (via 5 μL of 0.01 N HCl), repeated 4–6 times. A plot of pH vs. added H^+^ was fitted by linear regression. The slope (γ) is negative and smaller in magnitude for samples with higher buffer capacity. Thus, buffer capacity is defined as |γ|^−1^, and corrected transport activity was expressed as α × |γ|^−1^.

To compare wild-type and mutant SyHR, expression levels were estimated by the magnitude of flash-induced absorbance changes at 540 nm, near the dark-state absorption maximum. After transport measurements, cells were pelleted by centrifugation, resuspended in 50 mM citric acid buffer (pH 6) containing 0.2 M NaCl, and adjusted to OD_660_ = 10.0. Cells were lysed by sonication, and lysates were used to record flash-induced absorbance changes. Maximum negative deflections between 0.05–0.5 ms after flash were used as proxies for relative expression levels. To correct for expression differences, transport activities were divided by the respective expression level for each sample.

Anion transport activity was analyzed with respect to dehydrated volume and p*K*a. Dehydrated volumes of polyatomic ions were calculated using Winmostar V11 (X-Ability Co. Ltd., Japan) in van der Waals molecular surface mode. Spherical volumes of Cl^−^, Br^−^, and I^−^ were calculated based on their previously reported ionic radii ^27^. p*K*a values were calculated using MarvinSketch 25.1.3 (Chemaxon, https://www.chemaxon.com).

### Absorption spectrum and flash photolysis spectroscopies

Lipid-reconstituted SyHR readily precipitated in salt-containing solutions and was therefore unsuitable for direct spectral measurements. Instead, the protein was resuspended in a salt-free buffer and embedded in 15% acrylamide gels, as previously described ^28^. For each tested inorganic or organic anion, 10 to 15 gels were prepared. Buffer exchange was performed by soaking the gels in 50 mM citric acid–NaOH buffer (pH 6) containing varying concentrations of the target anion, added as its sodium salt. Sodium gluconate was also included to maintain ionic strength. Due to its limited solubility, the maximum sodium gluconate concentration used was 1 M— equivalent in ionic strength to 1 M monovalent salt. For methanesulfonate and ethanesulfonate, sodium gluconate was included at concentrations up to 1 M, but it was omitted at higher anion concentrations.

Absorption spectra were recorded at room temperature using a UV1800 spectrometer (Shimadzu). Flash-induced absorbance changes were measured using a single-wavelength kinetic spectrometer, as described previously ^29-30^. Actinic excitation was provided by the second harmonic of a Q-switched Nd:YAG laser (532 nm, 5 ns, 1 mJ per pulse). Transient absorbance changes at selected wavelengths were recorded and averaged over 20 laser pulses to improve the signalto-noise ratio. All measurements were performed at 23 °C.

## RESULTS

### Demonstration of organic anion transport by ClRs

Chloride-pumping rhodopsins (ClRs) can be phylogenetically classified into three distinct groups (Fig. 1B) ^31^: archaeal ClRs^32-33^, marine eubacterial ClRs ^34^, and cyanobacterial ClRs ^20^.

We selected one or two representative members from each group to examine their ability to transport organic anions. Several ClRs are abbreviated as “HR” (e.g., SyHR), a term originating from “halorhodopsin,” the name given to the first ClR discovered in a halophilic archaeon ^32-33^.

All tested ClRs were heterologously expressed in *E. coli* membranes. This enabled straightforward detection of inward anion pump activity by monitoring light-induced pH increases in *E. coli* suspensions in the presence of a protonophore such as CCCP. In this setup, anion uptake by ClRs generates a negative membrane potential, which drives passive H^+^ influx and raises the extracellular pH. Conventional pH electrodes, which use 3.3 M KCl as an internal solution, risk contaminating samples with chloride ions. To avoid this, we used a custom electrochemical cell incorporating ITO electrodes (Fig. S1A) ^17^. These transparent electrodes respond rapidly to pH changes. In our setup, the *E. coli* suspension is separated from a reference salt solution by a dialysis membrane. Illumination causes a pH increase only in the *E. coli* compartment, leading to a voltage difference (ΔV) between the two electrodes. A typical light-induced ΔV trace for FR, a marine ClR, is shown in Fig. S1B. One second of illumination produces a negative ΔV, indicating a pH increase in the *E. coli* suspension. To improve the signal-to-noise ratio, measurements were averaged over 30–50 trials. All tested anions were used at concentrations of 200 mM.

Figure 1C displays the light-induced voltage changes (ΔV) for SyHR in the presence of various anions. Corresponding results for other ClRs are presented in Fig. S2. In the top row of Fig. 1C, robust pump activity was confirmed for typical inorganic anions: (a) Cl^−^, (b) Br^−^, (c) NO_3_^−^, and (d) I^−^. Next, we evaluated short-chain fatty acids (SCFAs), including (e) acetate, (f) propionate, (g) butyrate, and (h) lactate. However, no pump activity was detected for SyHR or the other ClRs (Fig. S2).

Although acetate (51.3 Å^3^) is close in size to iodide (47.7 Å^3^), its significantly higher p*K*a (4.5 vs. −9.0) may explain the lack of transport. Other SCFAs also exhibit relatively high p*K*a values (3.8 to 4.9), in contrast with those of the inorganic anions tested (−9.0 to −1.4). Figure 2A summarizes the dehydrated volumes and p*K*a values of all anions examined. Given the hydrophobic environment of microbial rhodopsin transmembrane regions, weak acids like SCFAs likely remain uncharged inside the protein and thus fail to interact strongly with it, precluding transport.

**Figure 2.**
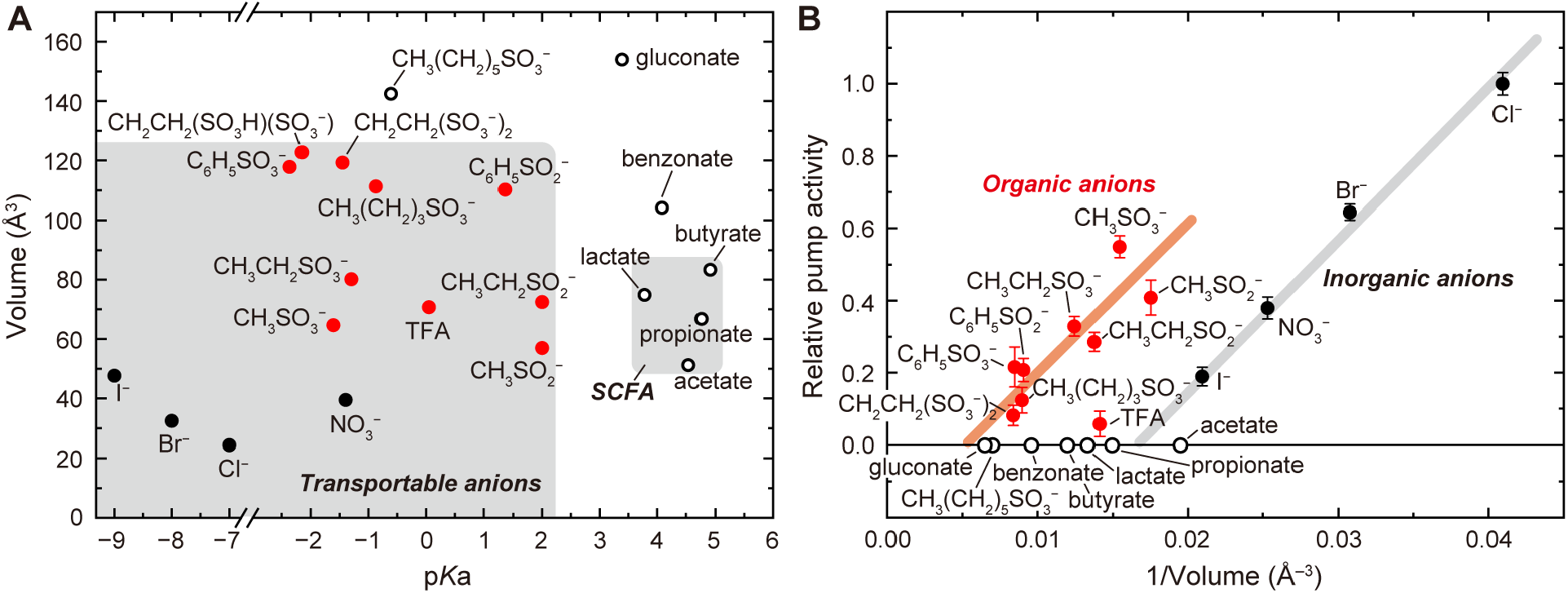
Dependence of anion transport activity on ion size and p*K*a. (A) p*K*a values and dehydrated volumes of the tested anions. Ethanedisulfonic acid has two p*K*a values; its lower and higher p*K*a are plotted in association with the dehydrated volumes of its monoanionic (CH_2_CH_2_(SO_3_H)(SO_3_ ^−^)_2_) and dianionic (CH_2_CH_2_(SO_3_ ^−^)_2_) forms, respectively. (B) Relationship between transport activity and the reciprocal of dehydrated ion volume. Error bars represent standard deviations (n = 30–50). The activity for ethanedisulfonate is plotted against the reciprocal volume of its dianionic form (CH_2_CH_2_(SO_3_ ^−^)_2_ ) In both panels, data for inorganic anions are shown as black filled circles, transportable organic anions as red filled circles, and non-transportable organic anions as black open circles. In (B), gray and orange diagonal lines are shown as visual guides to highlight distinct trends for inorganic and organic anions, respectively.

To test this hypothesis, we examined anions derived from stronger acids containing sulfinic (–SO_2_^−^) and sulfonic (–SO_3_^−^) groups, which have p*K*a values below 2 (Fig. 2A). In the third and fourth rows of Fig. 1C, distinct pump activities were observed for (i) methanesulfinate, (j) methanesulfonate, (k) ethanesulfinate, and (l) ethanesulfonate. Although transport efficiency decreased for larger anions (fourth row), SyHR still exhibited activity toward bulkier substrates, including (n) benzenesulfinate and (o) benzenesulfonate, both containing a benzene ring. In contrast, negligible activity was observed for (p) hexanesulfonate, likely due to its size (142.4 Å^3^) exceeding the upper limit for transport. We also tested additional anions with varying properties: (q) trifluoroacetate (TFA), (r) ethanedisulfonate, (s) benzoate, and (t) gluconate. Although TFA (70.8 Å^3^) is smaller than ethanedisulfonate (CH_2_CH_2_(SO _3_^−^)_2_ , 119.3 Å^3^), it exhibited lower transport activity, possibly due to its relatively high p*K*a of 0.05. In this study, “ethanedisulfonate” refers to the dianionic form; however, it remains uncertain whether the monoanionic or dianionic form is actually transported. Benzoate (104.2 Å^3^) is smaller than benzenesulfonate (C_6_H_5_SO_3_^−^, 117.9 Å^3^), but was not transported, likely due to its high p*K*a of 4.1. Gluconate, which has both a high p*K*a (3.4) and a large volume (153.9 Å^3^), was also not transported.

Pump activity for organic anions was also observed in two other ClRs: the archaeal NpHR and the marine eubacterial FR (Fig. S2). However, their activities were much weaker than that of SyHR, leading us to select SyHR for more detailed analysis.

As shown in Fig. 1C, SyHR exhibited the strongest activity with Cl^−^, while transport efficiency decreased for larger anions. However, the size dependence was not straightforward. For example, the pump activity for methanesulfonate (CH_3_SO_3_^−^) (j) was greater than that for NO_3_^−^ (c) and I^−^ (d), despite its larger molecular volume (64.7 Å^3^ vs 39.5 and 47.7 Å^3^) (Fig. 2A). To better understand this relationship, we plotted the relative pump activities against the reciprocal of ion volume (1/Volume) in Fig. 2B. The y-axis represents the initial slope of the light-induced voltage change, adjusted for the buffer capacity of each sample (see Materials and Methods and Fig. S3). The plot reveals a general trend: for both inorganic and organic anions, transport efficiency decreases as ion volume increases. However, this dependency differs between the two groups. Inorganic and organic anions follow distinct trends, illustrated by grey and orange guidelines, respectively. These results indicate that ion size alone cannot fully explain the differences in transport activity. Other factors—such as the ion’s chemical structure or interaction with the protein— appear to influence how effectively different anions are transported by SyHR.

### Organic anion binding to SyHR in the dark state

ClRs generally bind inorganic anions even in the dark state and subsequently transport them upon illumination ^20, 31, 34-35^. Figure 1A illustrates this initial binding state for SyHR, with Cl^−^ depicted as a green sphere. But do organic anions also bind in the dark? The binding site for inorganic ions is located near the protonated Schiff base (Fig. 1A), and their binding typically induces shifts in the absorption spectrum. These spectral shifts for inorganic anions are shown in Fig. 3A–D. In all cases, a blue shift was observed as the ion concentration increased. Figures 3E and 3F show the corresponding data for smaller sulfonate ions—methanesulfonate (CH_3_SO_3_^−^) and ethanesulfonate (CH_3_CH2SO_3_^−^), respectively. Distinct spectral changes occurred for both ions.

**Figure 3.**
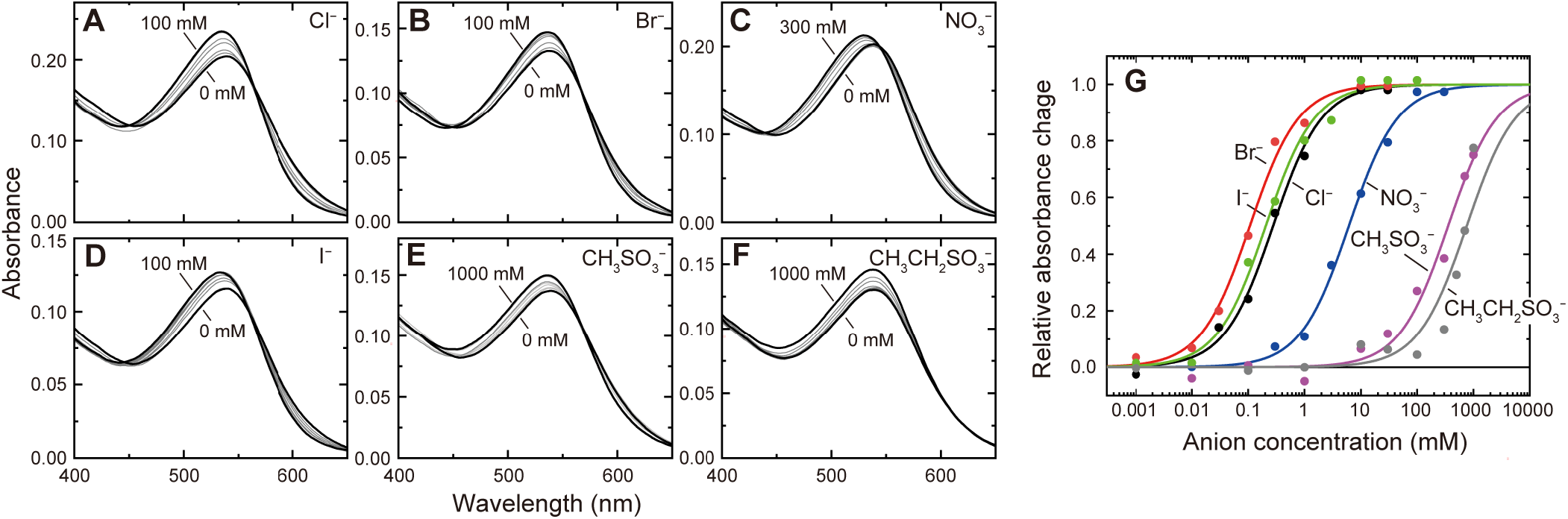
Anion binding to SyHR in the dark state. (A–F) Absorption spectral shifts of SyHR upon binding transportable anions. Lipid-reconstituted SyHR was used for all samples. Buffer solution contained 50 mM citric acid (pH 6) with various concentrations of sodium salts of the respective anions and sodium gluconate. Sodium gluconate was added to maintain constant ionic strength. (G) Absorbance changes at the wavelengths showing maximal responses were plotted against anion concentration and fitted with the Hill equation (n = 1) to determine dissociation constants, which are listed in Table S1. Relative absorbance changes are shown, along with best-fit curves.

To estimate the dissociation constants (*K*_d_), absorbance changes at wavelengths showing maximum response were plotted against ion concentration (Fig. 3G). The resulting *K*_d_ values are listed in Table S1. For inorganic anions, *K*_d_ values ranged from 0.11 to 6.15 mM, indicating that nearly all SyHR molecules were in the ion-bound state at 200 mM—the anion concentration used for transport activity measurements (Fig. 1C). For methanesulfonate and ethanesulfonate, saturation was not clearly observed (Fig. 3G), but we tentatively estimated *K*_d_ values of 356 mM and 761 mM, respectively. This suggests that at 200 mM, only about 36% and 21% of SyHR molecules were in the ion-bound state for methanesulfonate and ethanesulfonate, respectively. Although the affinities are significantly weaker, both organic anions can indeed bind to SyHR in the dark.

### Mutation effects of the residues constituting the Cl^−^ binding site

Do organic anions share the same binding site as inorganic ones? As shown in Fig. 1A, the binding of inorganic ions is stabilized by hydrogen bonds with Thr74 and Ser78. If both types of ions bind at the same site, mutations at these residues would be expected to reduce transport activity for both groups. Figure S4 shows the light-induced voltage changes detected by ITO electrodes for the T74A and S78A mutants. In Fig. 4, the initial slopes of these traces— representing transport activity—are plotted as open circles. For comparison, the corresponding data for wild-type SyHR (reproduced from Fig. 2B) are shown as filled circles. Mutant data were corrected for differences in protein expression levels relative to wild-type (see Materials and Methods and Fig. S5). As shown in Fig. 4A and 4B, both mutants exhibited clearly reduced transport activity for both inorganic and organic anions. This suggests that both ion types share the same binding site. However, transport activity for organic anions was nearly abolished in both mutants, indicating that the functional importance of these residues differs between the two groups. Inorganic anions can still bind and be transported in the absence of either residue, albeit less efficiently. In contrast, both residues appear essential for stable binding of organic anions.

**Figure 4.**
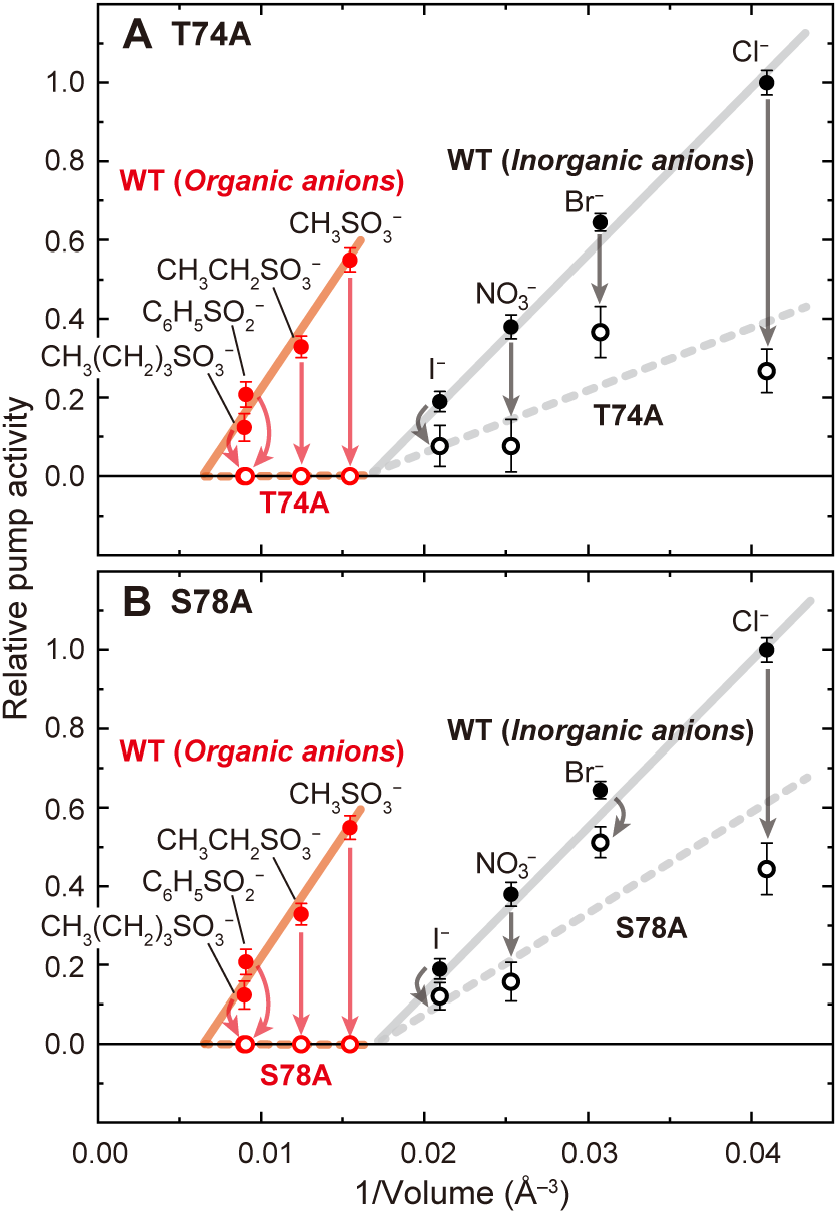
Reduced anion transport activity in SyHR mutants. Data for the T74A (A) and S78A (B) mutants are shown as open circles. For comparison, the corresponding data for wild-type SyHR are shown as filled circles (identical to those in Fig. 2B). Error bars represent standard deviations (n = 30–50). Mutant data were corrected for differences in expression levels relative to wild-type SyHR.

### Photocycles of Cl^−^ vs. organic ion transport

All ClRs undergo photocycles even in the absence of transportable anions. However, the details differ from those of active ion-pumping cycles, reflecting ion movement within the protein. This distinction is also expected to apply to SyHR when comparing photocycles with and without organic ion transport. Figures 5A–C illustrate these differences. We plotted flash-induced absorbance changes at three representative wavelengths: in the absence of transportable ions (black lines in all panels), and in the presence of Cl^−^ (Fig. 5A), methanesulfonate (Fig. 5B), and ethanesulfonate (Fig. 5C). SyHR concentrations were kept constant across all samples. Representative light-minus-dark spectra are shown in Fig. S6.

**Figure 5.**
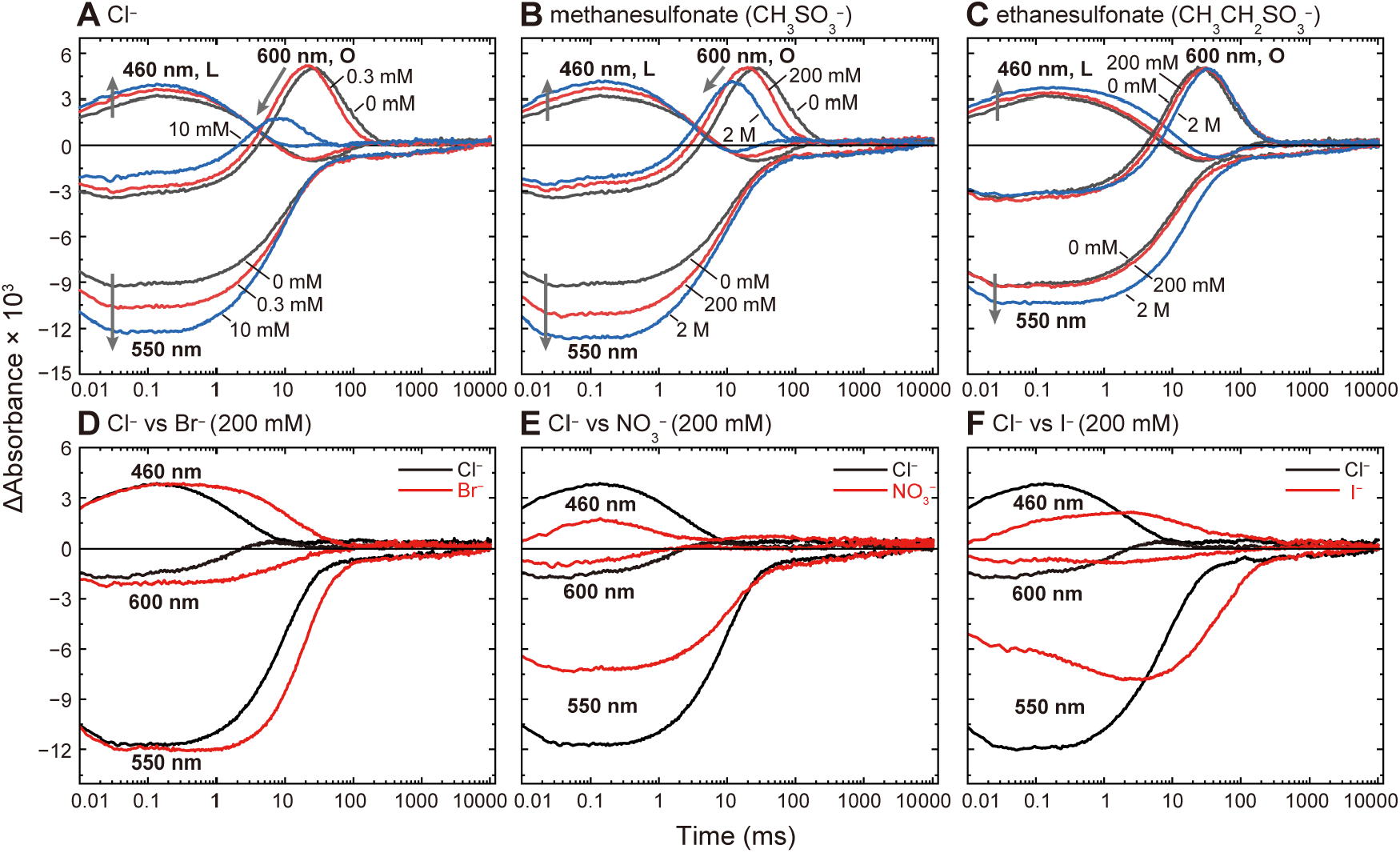
Photocycles of SyHR. Flash-induced absorbance changes at three representative wavelengths are plotted in each panel. SyHR concentration was kept constant across all samples. (A–C) Comparison of photocycles in the absence (black traces) and presence of (A) Cl^−^, (B) methanesulfonate, and (C) ethanesulfonate. (D–F) Comparison of the Cl^−^ transport photocycle with those of (D) Br^−^, (E) NO_3_^−^, and (F) I^−^. Sample and buffer conditions were essentially the same as those used in Fig. 3.

As discussed below, the photocycles associated with methanesulfonate and ethanesulfonate transport are essentially the same as that of Cl^−^, and all three differ from the photocycle observed without transportable anions. To ensure sufficient binding, we increased the concentrations of methanesulfonate and ethanesulfonate up to 2 M. At these highest concentrations, ∼85% and ∼72% of SyHR molecules are estimated to be bound, respectively, based on their dissociation constants (Table S1).

In the dark state, SyHR has a maximum absorbance at 530– 540 nm regardless of whether Cl^−^ or organic anions are bound (Fig. 3A, E, F). Accordingly, in Fig. 5A–C, the 550 nm traces exhibit early negative deflections, reflecting depletion of the ground state. As the concentration of Cl^−^ or organic anions increases, the magnitude of this deflection also increases, likely reflecting higher quantum efficiency of the substrate-bound form initiating the photocycle.

Positive absorbance changes at 460 nm and 600 nm correspond to the formation of two major intermediates common to Fig. 5A–C. Based on previous studies, the earlier intermediate at 460 nm is assigned to the L state, and the later one at 600 nm to the O state ^16^. As substrate concentration increases, the 460 nm signal becomes larger, indicating greater L formation. The O intermediate also shows clear substrate dependence under Cl^−^ and methanesulfonate conditions (Fig. 5A, B). As their concentrations increase, O accumulation decreases, suggesting that O is the substrate-releasing state and forms a quasi-equilibrium with L—i.e., L still retains the substrate. The decay of O also accelerates at higher concentrations of Cl^−^ and methanesulfonate, likely reflecting re-binding of the substrate from the extracellular side.

In contrast, ethanesulfonate had little effect on O accumulation, even at 2 M (Fig. 5C). Its larger size may weaken binding on the cytoplasmic side, shifting the L–O equilibrium toward O. Its size may also hinder uptake from the extracellular side, resulting in no measurable acceleration of O decay.

Following O decay in all three conditions (Fig. 5A–C), a small negative signal persists at 550 nm. This signal corresponds to the slow decay of the SyHR′ intermediate, a precursor of the ground state. The decay rate of SyHR′ was independent of substrate concentration and consistent across all three photocycles.

### Photocycles of inorganic anion transport

At 200 mM, essentially all SyHR molecules are in the ion-bound state for inorganic anions (Fig. 3G). Nonetheless, SyHR transports larger inorganic anions less efficiently (Figs. 1C and 2B). This difference in activity likely reflects differences in their respective photocycles. In Fig. 5D–F, flash-induced absorbance changes for 200 mM Cl^−^ are compared with those for 200 mM Br^−^, NO_3_^−^, and I^−^, respectively. SyHR concentrations were kept constant across all samples. Corresponding difference spectra are shown in Fig. S7.

As shown in Fig. 5D, the photocycle for Br^−^ is essentially the same as for Cl^−^. However, L decay is clearly slower for Br^−^. In contrast, distorted photocycles were observed in the presence of NO_3_^−^ and I^−^ (Figs. 5E, F). In both cases, the signals at 550 and 460 nm were markedly smaller, indicating a reduced fraction of SyHR molecules undergoing a productive photocycle. Furthermore, the L intermediate decayed in two steps, with the second decay occurring over several hundred milliseconds to seconds. These altered photocycles likely underlie the reduced transport activity observed for larger inorganic anions.

## DISCUSSION

We demonstrated that several ClRs can transport organic anions, with SyHR being particularly effective. SyHR was able to transport anions with molecular volumes up to 119.3 Å^3^ (e.g., ethanedisulfonate, CH_2_CH_2_(SO_3_^−^)_2_). All transported anions had p*K*a values below 2. Anions with higher p*K*a values were not transported, even if their volumes were smaller than 119.3 Å^3^. Thus, the ability to retain a negative charge within the protein appears to be a prerequisite for transport.

Like inorganic ions, certain organic anions—at least methanesulfonate and ethanesulfonate—can bind to unphotolyzed SyHR. Mutations at residues forming the Cl^−^ binding site impaired the transport of both inorganic and organic anions, indicating that they share the same binding site and that initial binding is essential for transport. This binding may require a low p*K*a, as the local pH near the protonated Schiff base may be close to 2—implying that anions with p*K*a values above 2 would remain uncharged and therefore unable to bind, even if their molecular volumes are small.

For both inorganic and organic anions, SyHR appears to transport only those that can bind to the protein in the dark state. Therefore, transport efficiency seems to be determined by two main factors: (1) the proportion of SyHR molecules in the ion-bound dark state, and (2) the efficiency of the transport reaction following photoactivation. For inorganic ions, nearly all SyHR molecules are in the bound state at 200 mM. However, larger inorganic ions trigger inefficient photoreactions, characterized by reduced photocycling and distorted photocycles. These features likely account for their lower transport activity. In contrast, SyHR molecules bound to organic anions undergo productive photoreactions. Although we were able to analyze the photocycles only of methanesulfonate and ethanesulfonate due to their relatively weak binding, both showed photocycles essentially identical to that of Cl^−^. While O decay was slower for organic anions, the subsequent decay of the SyHR′ intermediate—the final step of the photocycle— occurred at rates comparable to those in Cl^−^ transport. This suggests that the efficiency of the transport reaction itself is similar. Therefore, for organic anions, the reduced transport activity of larger ions is primarily due to the smaller proportion of SyHR molecules in the ion-bound state prior to illumination.

The strong binding of inorganic anions implies a high degree of similarity in size and/or shape to Cl^−^. Despite this similarity, SyHR exhibits inefficient photocycles when bound to larger inorganic anions. In contrast, organic anions bind only weakly to the unphotolyzed SyHR, yet they can still initiate productive photoreactions. Thus, strong binding does not necessarily guarantee a productive photoreaction. The detailed relationship between initial binding and subsequent photoreaction efficiency warrants further investigation.

In addition to SyHR, we found that NpHR and FR possess the ability to transport organic anions. This suggests that ClRs capable of transporting organic ions may be more widespread than we initially expected. Expanding the search may uncover ClRs capable of transporting even larger organic anions. SyHR and FR are closely related to the ClRs from *Mastigocladopsis repens* (MrHR) and *Nonlabens marinus* S1-08 (NM-R3), respectively, with amino acid sequence identities of 80.3% and 54.2% ^15-16, 20, 34^. However, MrHR and NM-R3 were unable to transport even smaller organic anions (Fig. S8), suggesting that subtle differences in residues and/or structural features determine the ability to transport organic anions. Identifying these determinants and exploring how to extend substrate range represent important directions for future investigation.

## ASSOCIATED CONTENT

### Supporting Information file includes

Photochemical cell for measuring anion transport activity (Fig. S1), Anion transport activities of ClRs other than SyHR (Fig. S2), Relative buffer capacities of samples used in anion transport activity measurements (Fig. S3), Anion transport activities of SyHR mutants (Fig. S4), Relative expression levels of wild-type SyHR and its mutants in *E. coli* membranes (Fig. S5), Photocycles of SyHR in the absence and presence of Cl^−^, methanesulfonate, and ethanesulfonate (Fig. S6), Photocycles of SyHR in the presence of 200 mM inorganic anions (Fig. S7), Anion transport activities of MrHR and NM-R3 (Fig. S8), Dissociation constants of SyHR for various anions (Table S1), References (PDF)

## ACKNOWLEDGMENT

This research was supported by JSPS KAKENHI Grant Numbers JP22H02579 (to T.T. and T.K.) and JP22H05389 (to T.K.); the Steel Foundation for Environmental Protection Technology Grant Number C-40-56 (to T.K.); and the JKA Foundation Grant Number 2025M-319 (to T.K.). Additional support was provided by the G-7 Scholarship Foundation Grant Number 7102400053 (to Y.S.).

## Notes

### Competing Interest Statement

The authors have declared no competing interest.

## REFERENCES

(1) Ernst, O. P.; Lodowski, D. T.; Elstner, M.; Hegemann, P.; Brown, L. S.; Kandori, H., Microbial and animal rhodopsins: structures, functions, and molecular mechanisms. Chem. Rev. 2014, 114 (1), 126–163.

(2) Govorunova, E. G.; Sineshchekov, O. A.; Li, H.; Spudich, J. L., Microbial Rhodopsins: Diversity, Mechanisms, and Optogenetic Applications. Annu. Rev. Biochem. 2017, 86, 845–872.

(3) Kandori, H., Biophysics of rhodopsins and optogenetics. Biophys. Rev. 2020, 12 (2), 355–361.

(4) Rozenberg, A.; Inoue, K.; Kandori, H.; Béjà, O., Microbial Rhodopsins: The Last Two Decades. Annu. Rev. Microbiol. 2021, 75, 427–447.

(5) Oesterhelt, D.; Stoeckenius, W., Rhodopsin-like protein from the purple membrane of Halobacterium halobium. Nature 1971, 233 (39), 149–152.

(6) Oesterhelt, D.; Stoeckenius, W., Functions of a new photoreceptor membrane. Proc. Natl. Acad. Sci. USA 1973, 70 (10), 2853–2857.

(7) Deisseroth, K., Optogenetics: 10 years of microbial opsins in neuroscience. Nat. Neurosci. 2015, 18 (9), 1213–25.

(8) Vlasova, A. D.; Bukhalovich, S. M.; Bagaeva, D. F.; Polyakova, A. P.; Ilyinsky, N. S.; Nesterov, S. V.; Tsybrov, F. M.; Bogorodskiy, A. O.; Zinovev, E. V.; Mikhailov, A. E.; Vlasov, A. V.; Kuklin, A. I.; Borshchevskiy, V. I.; Bamberg, E.; Uversky, V. N.; Gordeliy, V. I., Intracellular microbial rhodopsin-based optogenetics to control metabolism and cell signaling. Chem. Soc. Rev. 2024, 53 (7), 3327–3349.

(9) Deisseroth, K.; Hegemann, P., The form and function of channelrhodopsin. Science 2017, 357 (6356), eaan5544.

(10) Govorunova, E. G.; Sineshchekov, O. A.; Janz, R.; Liu, X.; Spudich, J. L., NEUROSCIENCE. Natural light-gated anion channels: A family of microbial rhodopsins for advanced optogenetics. Science 2015, 349 (6248), 647–50.

(11) Kojima, K.; Shibukawa, A.; Sudo, Y., The Unlimited Potential of Microbial Rhodopsins as Optical Tools. Biochemistry 2020, 59 (3), 218–229.

(12) Nakao, S.; Kojima, K.; Sudo, Y., Phototriggered Apoptotic Cell Death (PTA) Using the Light-Driven Outward Proton Pump Rhodopsin Archaerhodopsin-3. J. Am. Chem. Soc. 2022, 144 (9), 3771–3775.

(13) Duschl, A.; Lanyi, J. K.; Zimányi, L., Properties and photochemistry of a halorhodopsin from the haloalkalophile, Natronobacterium pharaonis. J. Biol. Chem. 1990, 265 (3), 1261–1267.

(14) Seki, A.; Miyauchi, S.; Hayashi, S.; Kikukawa, T.; Kubo, M.; Demura, M.; Ganapathy, V.; Kamo, N., Heterologous expression of pharaonis halorhodopsin in Xenopus laevis oocytes and electrophysiological characterization of its light-driven Cl^-^pump activity. Biophys. J. 2007, 92 (7), 2559–2569.

(15) Inoue, K.; Koua, F. H.; Kato, Y.; Abe-Yoshizumi, R.; Kandori, H., Spectroscopic study of a light-driven chloride ion pump from marine bacteria. J. Phys. Chem. B 2014, 118 (38), 11190–9.

(16) Niho, A.; Yoshizawa, S.; Tsukamoto, T.; Kurihara, M.; Tahara, S.; Nakajima, Y.; Mizuno, M.; Kuramochi, H.; Tahara, T.; Mizutani, Y.; Sudo, Y., Demonstration of a light-driven SO_4_^2-^transporter and its spectroscopic characteristics. J. Am. Chem. Soc. 2017, 139, 4376–4389.

(17) Doi, Y.; Watanabe, J.; Nii, R.; Tsukamoto, T.; Demura, M.; Sudo, Y.; Kikukawa, T., Mutations conferring SO_4_^2−^ pumping ability on the cyanobacterial anion pump rhodopsin and the resultant unique features of the mutant. Sci. Rep. 2022, 12 (1), 16422.

(18) Singh, M.; Ito, S.; Hososhima, S.; Abe-Yoshizumi, R.; Tsunoda, S. P.; Inoue, K.; Kandori, H., Light-Driven Chloride and Sulfate Pump with Two Different Transport Modes. J. Phys. Chem. B 2023, 127 (32), 7123–7134.

(19) Sato, M.; Kubo, M.; Aizawa, T.; Kamo, N.; Kikukawa, T.; Nitta, K.; Demura, M., Role of putative anion-binding sites in cytoplasmic and extracellular channels of Natronomonas pharaonis halorhodopsin. Biochemistry 2005, 44 (12), 4775–4784.

(20) Hasemi, T.; Kikukawa, T.; Kamo, N.; Demura, M., Characterization of a Cyanobacterial Chloride-pumping Rhodopsin and Its Conversion into a Proton Pump. J. Biol. Chem. 2016, 291 (1), 355–62.

(21) Lee, K. A.; Lee, S. S.; Kim, S. Y.; Choi, A. R.; Lee, J. H.; Jung, K. H., Mistic-fused expression of algal rhodopsins in Escherichia coli and its photochemical properties. Biochim. Biophys. Acta 2015, 1850 (9), 1694–703.

(22) Iizuka, A.; Kajimoto, K.; Fujisawa, T.; Tsukamoto, T.; Aizawa, T.; Kamo, N.; Jung, K. H.; Unno, M.; Demura, M.; Kikukawa, T., Functional importance of the oligomer formation of the cyanobacterial H^+^ pump Gloeobacter rhodopsin. Sci. Rep. 2019, 9 (1), 10711.

(23) Murabe, K.; Tsukamoto, T.; Aizawa, T.; Demura, M.; Kikukawa, T., Direct Detection of the Substrate Uptake and Release Reactions of the Light-Driven Sodium-Pump Rhodopsin. J. Am. Chem. Soc. 2020, 142 (37), 16023–16030.

(24) Sasaki, S.; Tamogami, J.; Nishiya, K.; Demura, M.; Kikukawa, T., Replaceability of Schiff base proton donors in light-driven proton pump rhodopsins. J. Biol. Chem. 2021, 297, 101013.

(25) Hamada, C.; Murabe, K.; Tsukamoto, T.; Kikukawa, T., Direct detection of the chloride release and uptake reactions of Natronomonas pharaonis halorhodopsin. J. Biol. Chem. 2024, 300 (9), 107712.

(26) Hasemi, T.; Kikukawa, T.; Watanabe, Y.; Aizawa, T.; Miyauchi, S.; Kamo, N.; Demura, M., Photochemical study of a cyanobacterial chloride-ion pumping rhodopsin. Biochim. Biophys. Acta Bioenerg. 2019, 1860 (2), 136–146.

(27) Marcus, Y., Ionic radii in aqueous solutions. Chem. Rev. 1988, 88, 1475–1498.

(28) Sato, Y.; Hashimoto, T.; Kato, K.; Okamura, A.; Hasegawa, K.; Shinone, T.; Tanaka, Y.; Tanaka, Y.; Tsukazaki, T.; Tsukamoto, T.; Demura, M.; Yao, M.; Kikukawa, T., Multistep conformational changes leading to the gate opening of light-driven sodium pump rhodopsin. J. Biol. Chem. 2023, 299 (12), 105393.

(29) Kikukawa, T.; Kusakabe, C.; Kokubo, A.; Tsukamoto, T.; Kamiya, M.; Aizawa, T.; Ihara, K.; Kamo, N.; Demura, M., Probing the Cl^-^-pumping photocycle of pharaonis halorhodopsin: Examinations with bacterioruberin, an intrinsic dye, and membrane potential-induced modulation of the photocycle. Biochim. Biophys. Acta Bioenerg. 2015, 1847 (8), 748–758.

(30) Kato, T.; Tsukamoto, T.; Demura, M.; Kikukawa, T., Real-time identification of two substrate-binding intermediates for the light-driven sodium pump rhodopsin. J. Biol. Chem. 2021, 296, 100792.

(31) Kikukawa, T., Functional Mechanism of Cl^−^-Pump Rhodopsin and Its Conversion into H^+^ Pump. In Optogenetics: Light-Sensing Proteins and Their Applications in Neuroscience and Beyond, Yawo, H.; Kandori, H.; Koizumi, A.; Kageyama, R., Eds. Springer Singapore: Singapore, 2021; pp 55–71.

(32) Matsuno-Yagi, A.; Mukohata, Y., Two possible roles of bacteriorhodopsin; a comparative study of strains of Halobacterium halobium differing in pigmentation. Biochem. Biophys. Res. Commun. 1977, 78 (1), 237–243.

(33) Schobert, B.; Lanyi, J. K., Halorhodopsin is a light-driven chloride pump. J. Biol. Chem. 1982, 257 (17), 10306–10313.

(34) Yoshizawa, S.; Kumagai, Y.; Kim, H.; Ogura, Y.; Hayashi, T.; Iwasaki, W.; DeLong, E. F.; Kogure, K., Functional characterization of flavobacteria rhodopsins reveals a unique class of light-driven chloride pump in bacteria. Proc. Natl. Acad. Sci. U.S.A. 2014, 111 (18), 6732–7.

(35) Váró, G.; Brown, L. S.; Sasaki, J.; Kandori, H.; Maeda, A.; Needleman, R.; Lanyi, J. K., Light-driven chloride ion transport by halorhodopsin from Natronobacterium pharaonis. I. The photochemical cycle. Biochemistry 1995, 34 (44), 14490–14499.

