## Supplementary material for "Discovery of Light-Powered Organic Ion Transport by a Natural Protein": Fig. S1; Fig. S2; Fig. S3; Fig. S4; Fig. S5; Fig. S6; Fig. S7; Fig. S8; Table S1

**Table of Contents**


Figure S1. Photochemical cell for measuring anion transport activity.................................S3

Figure S2. Anion transport activities of ClRs other than SyHR...........................................S4

Figure S3. Relative buffer capacities of samples used in anion transport activity measurements..............................................................S5

Figure S4. Anion transport activities of SyHR mutants.......................................................S6

Figure S5. Relative expression levels of wild-type SyHR and its mutants in *E. coli* membranes..........................................................................S7

Figure S6. Photocycles of SyHR in the absence and presence of $Cl^-$, methanesulfonate, and ethanesulfonate......................................................S8

Figure S7. Photocycles of SyHR in the presence of 200 mM inorganic anions..................S9

Figure S8. Anion transport activities of MrHR and NM-R3.............................................S10

Table S1. Dissociation constants of SyHR for various anions...........................................S11

References...........................................................................................................S11

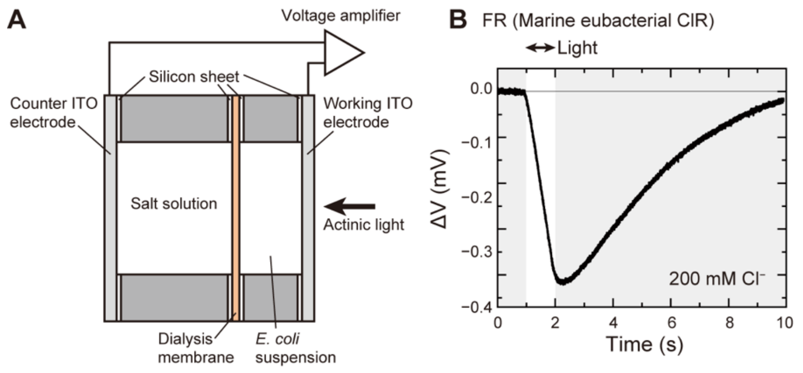


**Figure S1.** Photochemical cell for measuring anion transport activity. (A) Schematic illustration of the photochemical cell. Inward anion transport by a ClR induces a pH increase in the *E. coli* suspension. This change is detected as a voltage difference between two ITO electrodes. (B) Representative time course of a light-induced voltage change. Actinic light was applied from 1 to 2 seconds. The sample contained *E. coli* cells expressing FR, a marine eubacterial ClR, suspended in 200 mM NaCl solution with 10 µM CCCP. The illustration in (A) is adapted from the previous report [1], licensed under CC BY 4.0.

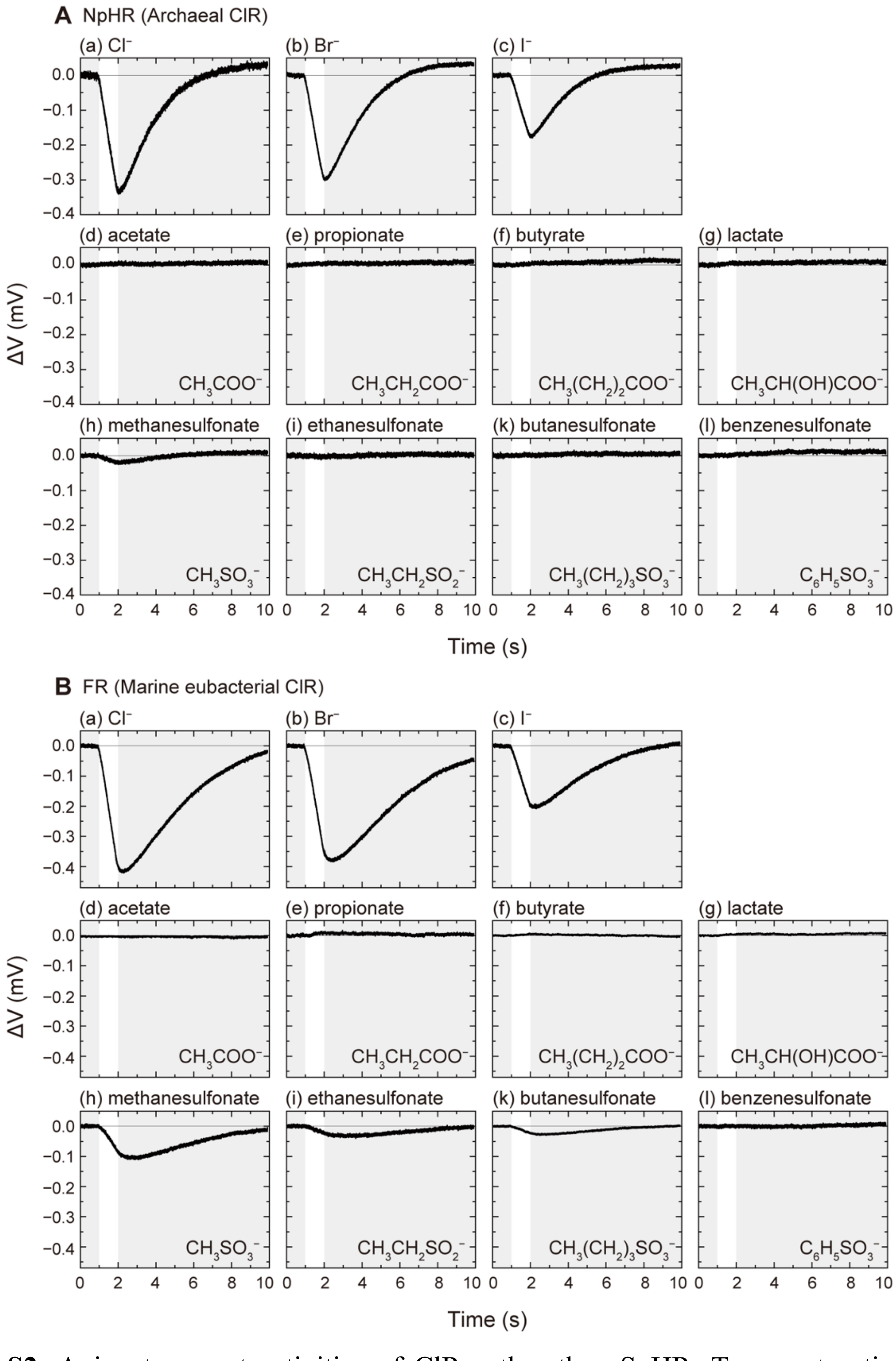


**Figure S2.** Anion transport activities of ClRs other than SyHR. Transport activity data corresponding to those shown in Fig. 1C are presented for (A) NpHR, an archaeal ClR, and (B) FR, a marine eubacterial ClR.

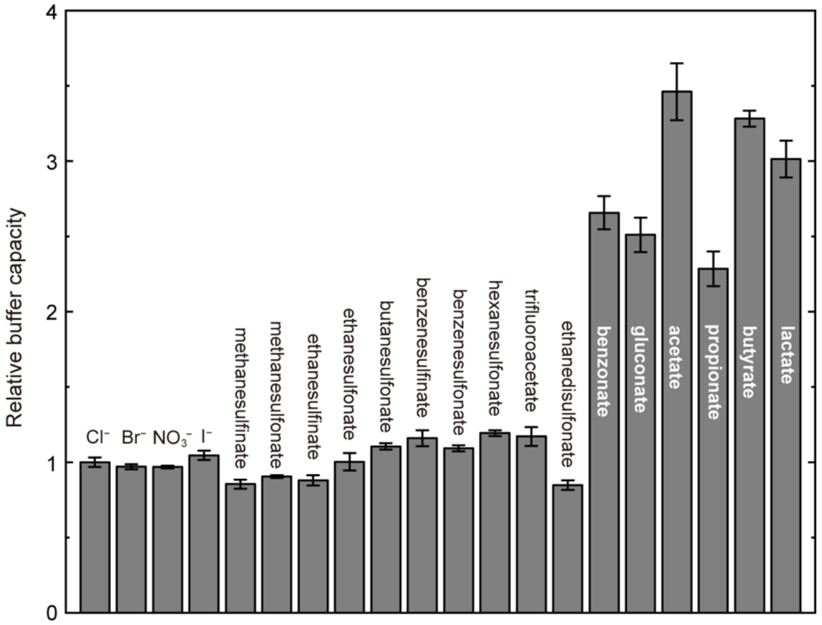


**Figure S3.** Relative buffer capacities of samples used in anion transport activity measurements. Error bars represent standard errors.

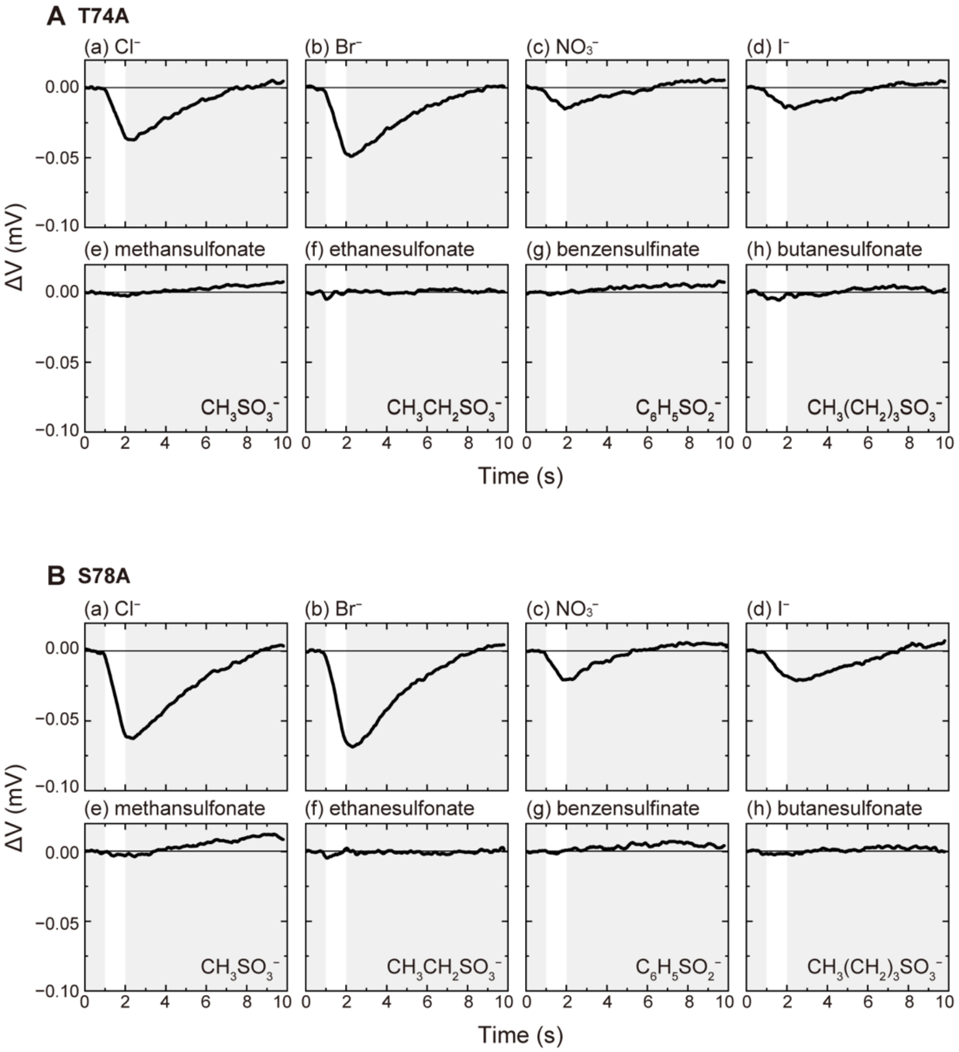


**Figure S4.** Anion transport activities of SyHR mutants. Data corresponding to those in Fig. 1C are shown for (A) T74A and (B) S78A SyHR mutants. The slopes of the light-induced voltage changes were determined by fitting analysis and used to generate the plots shown in Fig. 4.

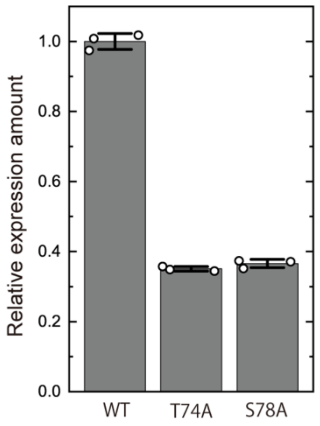


**Figure S5.** Relative expression levels of wild-type SyHR and its mutants in *E. coli* membranes. Expression levels were estimated from the maximum values of flash-induced absorbance changes at 540 nm. Open circles represent individual measurements, and bars indicate the mean ± standard deviation (n = 3).

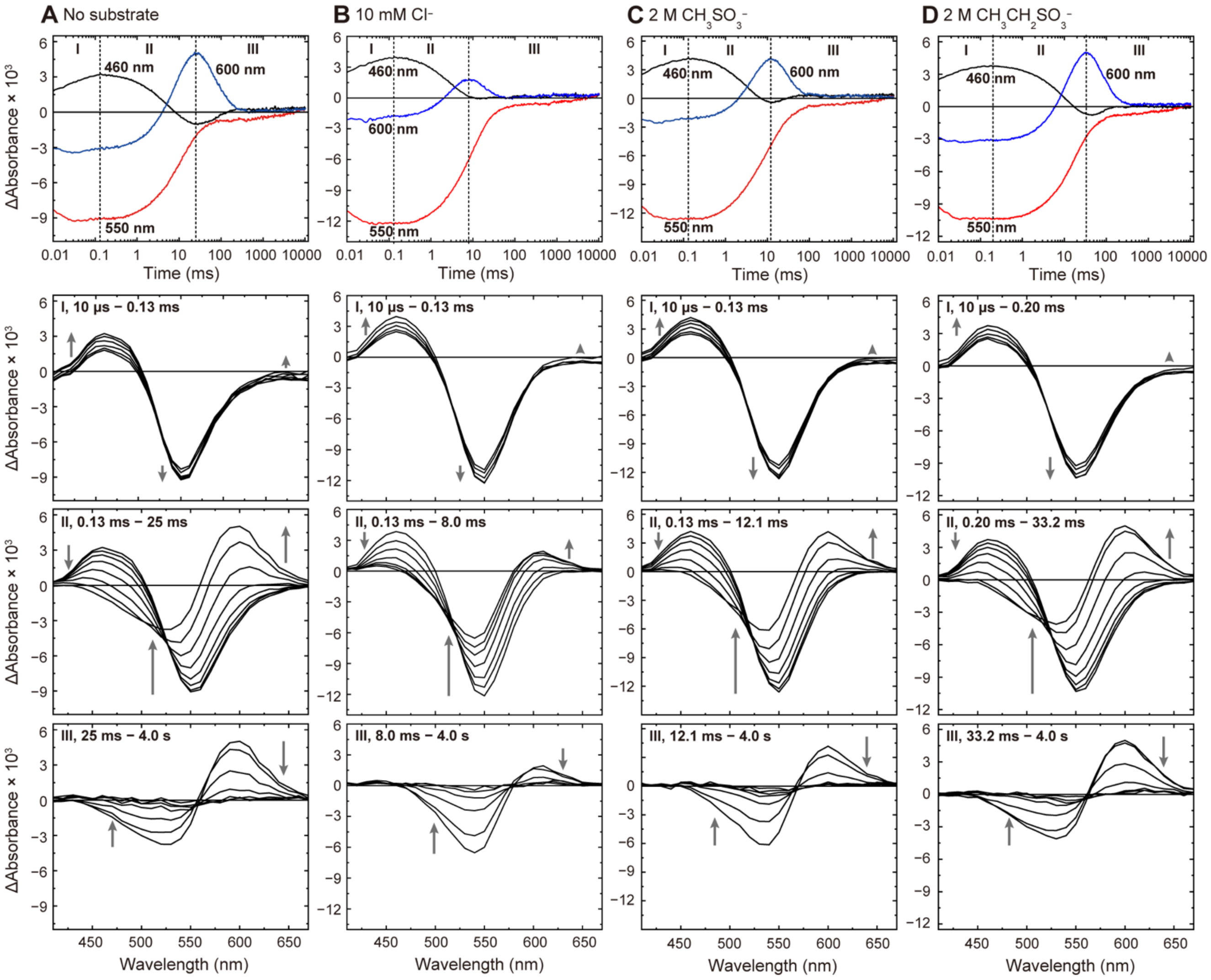


**Figure S6.** Photocycles of SyHR in the absence and presence of $Cl^-$, methanesulfonate, and ethanesulfonate. Top panels show flash-induced absorbance changes at three representative wavelengths (also shown in Fig. 5). Bottom panels show the corresponding light-minus-dark difference spectra, divided into three time regions (I–III), as indicated by vertical dashed lines in the top panels. SyHR reconstituted into lipid membranes was encapsulated in 15% acrylamide gels. These gels were immersed in 50 mM citric acid buffer (pH 6) containing sodium salts of the respective anions. For the samples in (A) no substrate and (B) 10 mM $Cl^-$, sodium gluconate was added to maintain ionic strength equivalent to 1 M of a monovalent salt. Sodium gluconate was not added to (C) 2 M methanesulfonate and (D) 2 M ethanesulfonate.

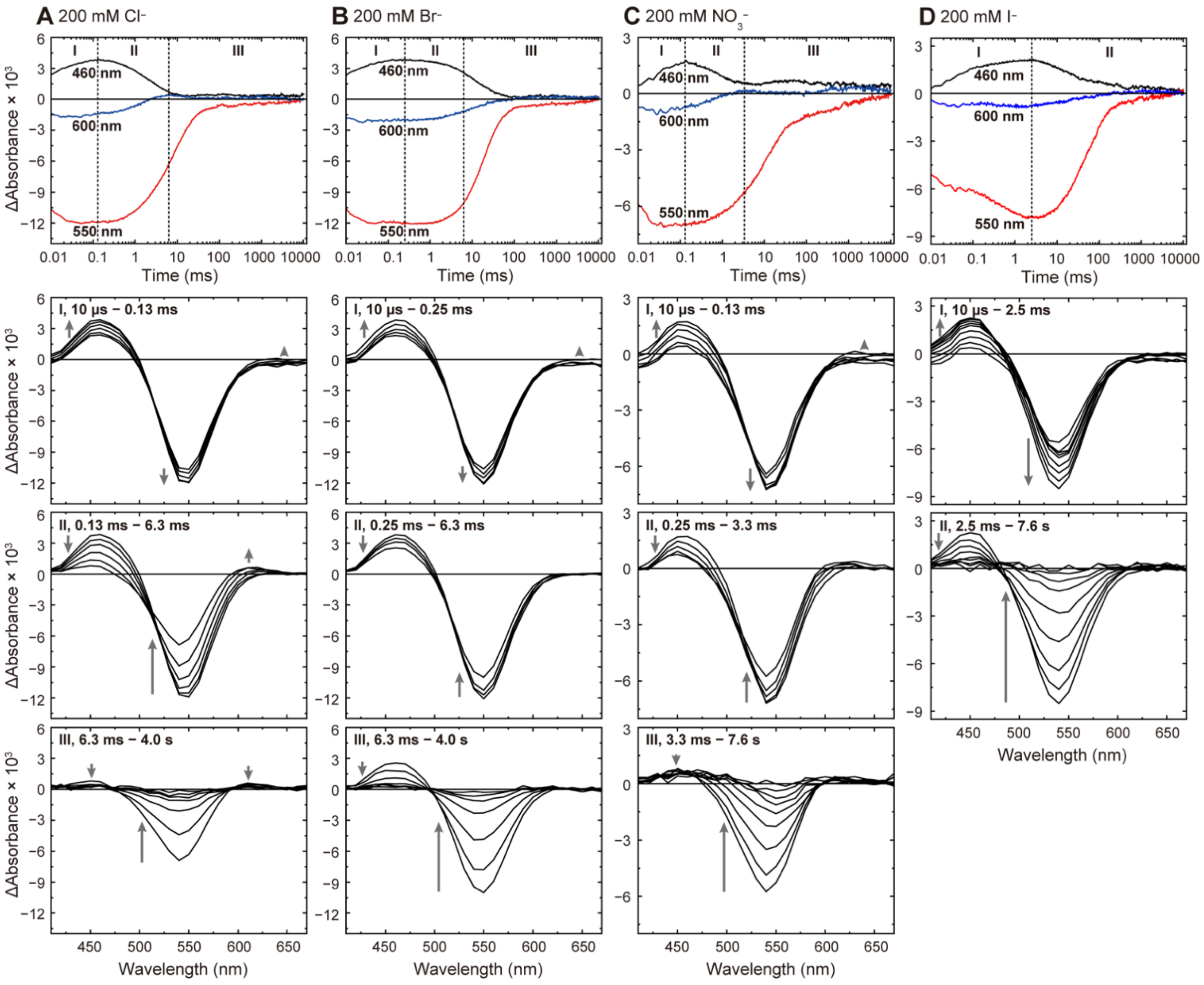


**Figure S7.** Photocycles of SyHR in the presence of 200 mM inorganic anions. Top panels show flash-induced absorbance changes at three representative wavelengths (also shown in Fig. 5). Bottom panels show the corresponding light-minus-dark difference spectra, divided into two or three time regions (I–II or I–III), as indicated by vertical dashed lines in the top panels. Sample conditions are the same as those described in Fig. 5.

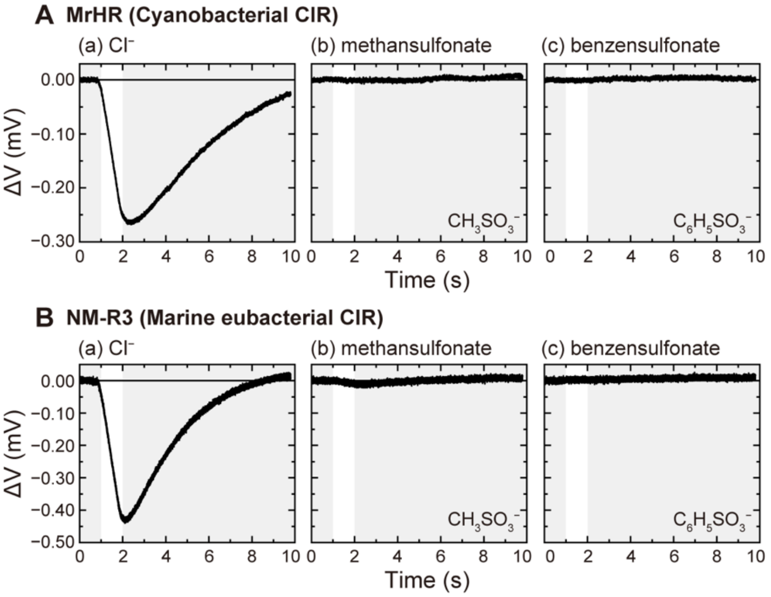


**Figure S8.** Anion transport activities of MrHR and NM-R3. Transport activity data are shown for (A) MrHR, a cyanobacterial ClR, and (B) NM-R3, a marine eubacterial ClR, with $Cl^-$, metansulfonate, and benzensulfonate as substrates. Sample conditions are the same as those described in Fig. 1C.

**Table S1.** Dissociation constants of SyHR for various anions.

| Anions | $K_d$ (mM) |
|---|---|
| $Cl^-$ | 0.28 ± 0.05 |
| $Br^-$ | 0.11 ± 0.02 |
| $NO_3^-$ | 6.15 ± 0.75 |
| $I^-$ | 0.21 ± 0.03 |
| $CH_3SO_3^-$ | 356 ± 133 |
| $CH_3CH_2SO_3^-$ | 761 ± 149 |

Values represent dissociation constants ($K_d$) determined by fitting absorbance changes to a binding curve using the Hill equation (n = 1, see Fig. 3G). Errors indicate standard errors of the fit.

### References

(1) Doi, Y.; Watanabe, J.; Nii, R.; Tsukamoto, T.; Demura, M.; Sudo, Y.; Kikukawa, T. Mutations conferring $SO_4^{2-}$ pumping ability on the cyanobacterial anion pump rhodopsin and the resultant unique features of the mutant. *Sci. Rep.* **2022**, *12* (1), 16422.
